# Differentiation of Peritubular Myoid-Like Cells from Human Induced Pluripotent Stem Cells

**DOI:** 10.1101/2021.06.04.447123

**Authors:** Meghan Robinson, Luke Witherspoon, Stephanie Willerth, Ryan Flannigan

**Affiliations:** Vancouver Prostate Centre, Vancouver, British Columbia; Department of Urologic Sciences, University of British Columbia, Vancouver, British Columbia, Canada; Department of Urology, The Ottawa Hospital, Ottawa, Ontario, Canada; Division of Medical Sciences, University of Victoria, Victoria, British Columbia, Canada; Department of Mechanical Engineering, University of Victoria, Victoria, British Columbia, Canada; School of Biomedical Engineering, University of British Columbia, Vancouver, British Columbia, Canada; Department of Urology, Weill Cornell Medicine, New York, NY, USA

**Author notes:** CORRESPONDING AUTHOR: Ryan Flannigan, MD, Department of Urologic Sciences, University of British Columbia, Vancouver, British Columbia, Gordon & Leslie Diamond Health Care Centre, Level 6, 2775 Laurel Street, Vancouver, BC Canada V5Z 1M9.

**Keywords:** Human Induced Pluripotent Stem Cells, Peritubular Myoid Cell, Leydig Cell, Cell Differentiation, Infertility, Platelet-Derived Growth Factor, Cell Culture, Testicular Cancer

## Abstract

Spermatogenesis is a complex process requiring intricate cellular interactions between multiple cell types to produce viable sperm. Peritubular myoid cells (PTMs) are smooth muscle cells that line the seminiferous tubules and play a critical role in sperm production by providing mechanical support and molecular signaling factors. *In vitro* investigation of their contribution to spermatogenesis and their dysfunction in infertility is currently limited by the rare accessibility of human testicular tissue for research. Therefore, this study set forth to generate an alternative source of PTMs using human induced pluripotent stem cells (hiPSCs) - adult cells that have been reprogrammed into a pluripotent state, making them capable of indefinite expansion and the regeneration of any cell type in the body. PTMs and Leydig cells arise from a common progenitor, so we hypothesized that PTMs could be derived by modifying an existing differentiation protocol for Leydig cell differentiation from hiPSCs. These hiPSC-derived cells, or hPTMs, were characterized and compared to hiPSC-derived Leydig cells (hLCs) and human primary Sertoli cells as a negative control. Our findings show that the substitution of the molecular patterning factor Platelet-Derived Growth Factor Subunit B (PDGF-BB) for Platelet-Derived Growth Factor Subunit A (PDGF-AA) in a molecule-based differentiation protocol for deriving Leydig-like cells, is sufficient to derive peritubular myoid-like cells. This study describes a method for generating PTM-like cells from hiPSCs. These cells will allow for ongoing understanding of the cellular interactions required for normal spermatogenesis in an *in vitro* setting.

## INTRODUCTION

Infertility affects approximately 15% of couples, with male factors contributing to 50% of cases.(Jarvi, et al., 2015) Development of treatments is hampered due to our limited understanding of the underlying mechanisms leading to this condition. One cell population implicated in infertility are the peritubular myoid cells (PTMs). These cells, in combination with Sertoli cells, constitute a barrier surrounding the seminiferous tubules. A frequent finding on cellular histology of infertile males are defects in the architecture of this barrier, suggesting a dysregulation in its formation or regulation,(Welter, et al., 2013) and consequently the local spermatogenic microenvironment.(Mayerhofer, 2013) PTMs occupy the boundary separating the interstitial Leydig cell compartment from the germ cell tubular compartment, and this strategic anatomic position suggests involvement in the regulation of multiple testicular cell populations. Despite their importance, PTMs remain poorly characterized.

Most of our understanding of PTMs comes from rodent knock-out models and rodent cell culture studies. These *in vitro* and *in vivo* studies have shown that PTMs are responsive to androgen stimulation, as evidenced by increased secretion of Sertoli cell regulatory factors(Skinner and Fritz, 1985) and the SSC self-renewal factor GDNF,(Chen, et al., 2016) and are vital to Leydig cell development and function.(Norton and Skinner, 1989, Welsh, et al., 2012) However, the comparative simplicity of the tubular wall architecture in rodents infers that PTM functionality may not be comparable to human PTM functionality.(Mayerhofer, 2013) Therefore, researchers have recently turned to *in vitro* and biopsy studies of human PTMs to provide greater insight into their nature and regulation.(Mayerhofer, 2013) However, accessibility to testis biopsies for research is limited to specialized clinical laboratories.

Advancing our understanding of PTMs requires a reproducible and accessible cellular model. To this end, human induced pluripotent stem cells (hiPSCs) are an attractive option for generating PTMs. These are adult cells which have been reprogrammed into an embryonic-like state by introducing genes essential for pluripotency(Takahashi, et al., 2007). Like embryonic stem cells, they can divide indefinitely and differentiate into all of the specialized cell types of the body.(Liu, et al., 2020)

Peritubular myoid cells share a common progenitor cell type with Leydig cells known as Stem Leydig Cells (SLCs). SLCs are a mesenchymal stem cell type specific to testis tissue, whose lineage has not yet been traced in humans, but in mice are known to arise from a Wilm’s Tumor 1 (WT1)-expressing progenitor of the gonadal primordium which gives rise to both SLCs and Sertoli progenitor cells.(Guo, et al., 2020, Liu, et al., 2016) There is accumulating evidence that Sertoli cell-secreted Platelet-Derived Growth Factor-AA (PDGFAA) and Platelet-Derived Growth Factor-BB (PDGFBB) are important patterning factors delineating Leydig cell and PTM cell fates from SLCs. Mice PDGFAA Receptor (PDGFRA) knock-out models fail to produce adult Leydig cells despite exhibiting normal development of SLCs,(Gnessi, et al., 2000) while Sertoli cell Androgen Receptor (AR) knock-out models exhibit reduced PDGFAA production, correlating with a 40% loss of Leydig cell differentiation.(De Gendt, et al., 2005) In contrast, PDGFBB signaling inhibits Leydig cell differentiation,(Odeh, et al., 2014) and induces a functional phenotype in PTMs,(Chiarenza, et al., 2000) including contractility and secretion of extracellular matrix proteins.(Gnessi, et al., 1993)

Therefore, in this study we hypothesized that substituting PDGFBB in place of PDGFAA into an existing Leydig cell differentiation protocol from hiPSCs(Chen, et al., 2019) would be enough to promote their differentiation into PTMs. We use immunocytochemistry and reverse transcription quantitative polymerase chain reaction (RT-qPCR) to show that cells with PTM-like or Leydig-like cell phenotypes and gene expression profiles can be alternatively derived from hiPSCs through the actions of PDGFBB or PDGFAA signaling. This study provides a means of generating PTM-like cells from hiPSCs for *in vitro* studies.

## MATERIALS AND METHODS

### Ethical approval

Testis biopsy samples were obtained through the University of British Columbia Andrology Biobank, with informed consent for research (CREB approved protocol H18-03543). Experiments using hiPSCs in this study were not subject to ethics approval from the University of British Columbia Clinical Research Ethics Boards or Stem Cell Oversight Committee, since they were derived from somatic cells and not intended for transfer into humans or non-human animals.

### Culture of hiPSCs

The hiPSC line used was the 1-DL-01 line from WiCell.(WiCell, 2021) hiPSCs were expanded on Growth Factor Reduced Matrigel (Corning, 354230) in mTeSR™-Plus medium (STEMCELL Technologies, 100-0276), and passaged when 90% confluent using ReLeSR™ enzyme-free selective passaging reagent (STEMCELL Technologies, 05872) to maintain purity.

### Differentiation of Leydig cells from hiPSCs

Leydig cells were differentiated from hiPSCs as previously described with a modification (**Figure 1A**).(Chen, Li, Chen, Xi, Zhao, Ma, Xu, Han, Zhao, Ge and Guo, 2019)

**Figure 1.**
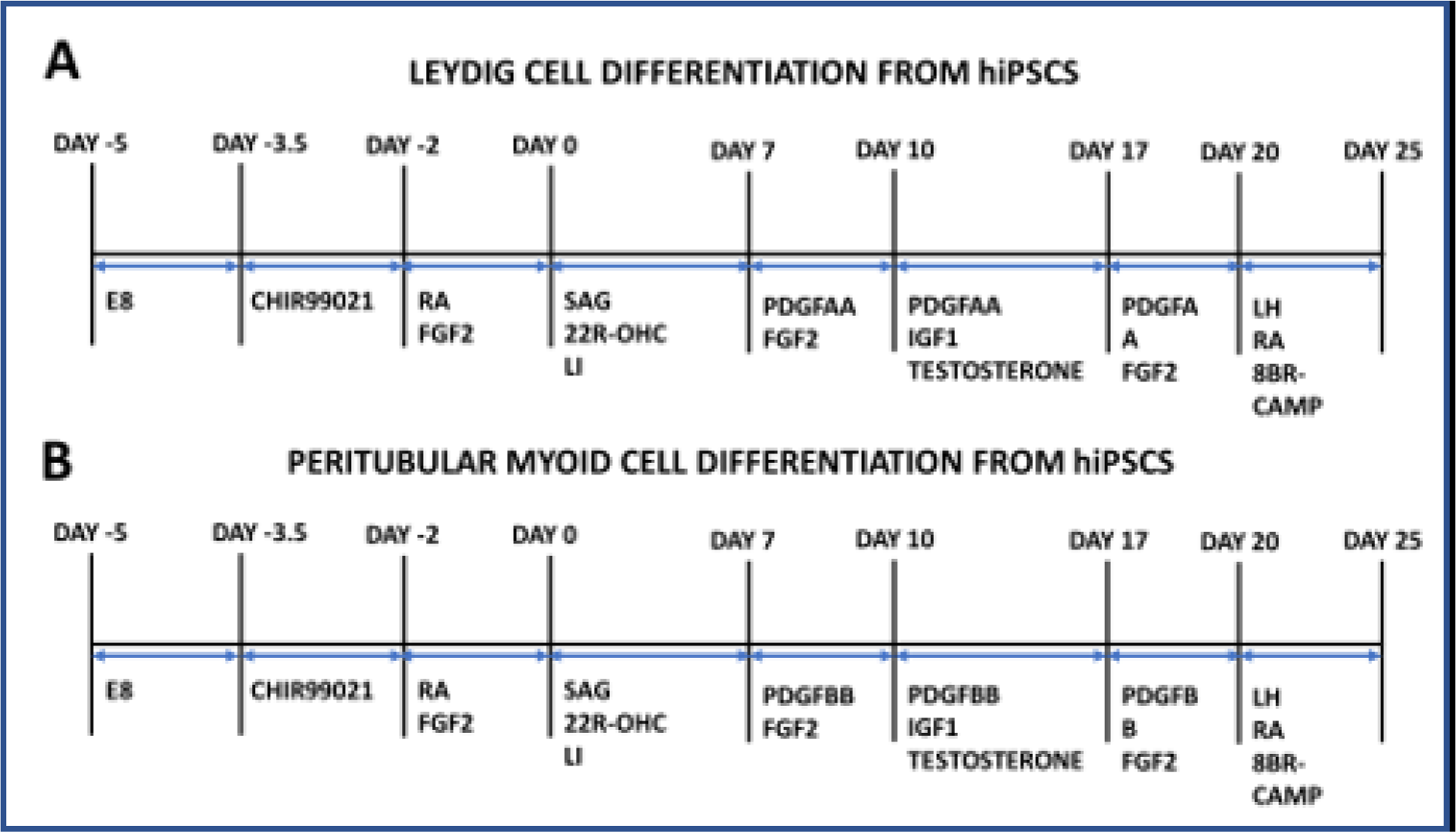
Timelines for differentiating Leydig cells and peritubular myoid cells from human induced pluripotent stem cells (hiPSCs). **A)** Timeline for differentiating Leydig cells from hiPSCs, based on a previously published protocol, with a modified preliminary step to first derive intermediate mesenchymal stem cells to improve differentiation efficiency. **B)** Timeline for differentiating peritubular myoid cells from hiPSCs by repeating the Leydig cell differentiation protocol, but substituting PDGFBB for PDGFAA. **Abbreviations:** E8 = TeSR-E8 medium, RA = retinoic acid, FGF2 = Fibroblast Growth Factor 2, SAG = Smoothened agonist, 22R-OHC = 22R-hydroxycholesterol, LI = lithium chloride, PDGFAA = Platelet-Derived Growth Factor-AA, PDGFBB = Platelet-Derived Growth Factor-BB, IGF1 = Insulin-Like Growth Factor 1, LH = luteinizing hormone, 8BR-CAMP = 8-bromo-cAMP.

The original protocol yielded 41.5% Leydig cells, likely because the hiPSCs spontaneously differentiated for 2 days prior to patterning, allowing for endodermal and ectodermal lineage acquisition in addition to mesenchymal. Therefore, we first patterned the hiPSCs with a mesenchymal protocol as previously described.(Lam, et al., 2014) Cells were treated with 5 µM CHIR99021 (STEMCELL Technologies, 72052) in Roswell Park Memorial Institute Medium 1640 (RPMI 1640, Gibco, 11875-093) for 36 hours, then 1 µM all-trans retinoic acid (RA, STEMCELL Technologies, 72262) and 100ng/mL human recombinant Fibroblast Growth Factor 2 (FGF2, STEMCELL Technologies, 78003.1) in RPMI 1640 for 2 days. They were then subjected to the following protocol for Leydig cell differentiation: cells were cultured throughout on Matrigel substrates in Dulbecco’s Modified Eagle Medium/Nutrient Mixture F-12 with 15mM HEPES (DMEM/F12, STEMCELL Technologies, 36254) 1X Insulin Transferrin Selenium Liquid Media Supplement (ITS, Millipore Sigma, I13146), 1% penicillin/streptomycin (Sigma Aldrich, P4333)and 1X GlutaMAX (Thermofisher, 35050079), 1% bovine serum albumin (BSA, Miltenyi Biotec, 130-091-376) and 5 ng/mL luteinizing hormone from human pituitary (LH, Sigma, L6420), with media changes every other day. From 0–7 days, 0.2 μM Smoothened Agonist (SAG, STEMCELL Technologies, 73412), 5 μM 22R-hydroxycholesterol (22R-OHC, Sigma, H9384), and 5 mM lithium chloride (Sigma, 62476) were added. From 7–10 days, 5 ng/mL human recombinant Platelet-Derived Growth Factor-AA (PDGF-AA, Peprotech, 100-13A) and 5 ng/mL FGF2 were added. From 10–17 days, 5 ng/mL PDGF-AA, 5 nM Insulin-Like Growth Factor 1 (IGF1, Peprotech, 100-11), and 10 μM testosterone (Toronto Research Chemicals, T155010) were added. From days 17–20, 10 ng/mL PDGF-AA and 10 ng/mL FGF2 were added. From days 20–25, 5 ng/mL LH (total of 10 ng/mL), 0.5 mM RA and 1 mM 8-bromo-cAMP (Peprotech, 2354843) were added. On day 25 cells the cells were switched to Leydig Cell Media (Sciencell Research Laboratories, 4511) for expansion, and passaged onto poly-L-lysine (PLL, Sciencell Research Laboratories, 0413) using TrypLE™ Express Enzyme (Thermofisher, 12604013).

### Differentiation of PTMS from hiPSCs

PTMs were differentiated from hiPSCs using the above protocol for hiPSC-derived Leydig cells with the modification that PDGF-AA is replaced by human recombinant Platelet-Derived Growth Factor-BB (PDGF-BB, Peprotech, 100-14B, **Figure 1B**). On day 25 the cells were expanded on PLL-coated plates in DMEM/F12, 1X ITS, 1% penicillin/streptomycin, 2.5% fetal bovine serum (FBS, Gibco, 12483-020), 10 ng/mL FGF2, 1 ng/mL human recombinant Leukemia Inhibitory Factor (LIF, STEMCELL Technologies, 78055.1) and 10 ng/mL human recombinant Epidermal Growth Factor (EGF, STEMCELL Technologies, 78006.1), and passaged using TrypLE™ Express Enzyme.

### Primary Sertoli cell isolation and culture

Testicular biopsies were transported to the lab on ice in Hypothermosol® FRS (STEMCELL Technologies, 07935). They were rinsed three times with Hank’s Balanced Salt Solution (HBSS, Millipore Sigma, 55021C), then cut into 1mm^3^ pieces with surgical scissors, and digested by Collagenase NB4 (Nordmark Biochemicals, S1745402) at 2PZU / 100 mg tissue, for 5 minutes at 37°C and 250 rpm. The sample was vigorously shaken and then incubated for a further 3 minutes at 37°C and 250 rpm. After spinning for 5 minutes at 200 x g, sedimented tubules were rinsed 3 times with HBSS, with spinning at 200 x g for 5 minutes in between each rinse. The sedimented tubules were re-suspended in 0.25% Trypsin/EDTA (Sigma, T3924) and 0.8 kU / 100mg DNase I (Sigma, D4263). The tubules were pipetted 3-5 times with a 5 mL pipette and incubated for 5 minutes at 37°C. This step was repeated for a total of 3 incubations. 10% FBS was added to stop the digestion, and the tubules were filtered through a 70 µm filter, then a 40µm filter, and centrifuged for 15 minutes at 600 x g. The cells were sorted into somatic and germ fractions by overnight plating in a single well on a tissue culture-treated 6-well plate. The next day, non-adherent cells were removed (germ cells) and the adherent cells (somatic cells) were expanded on PLL-coated plates in a 1:1:1 mixture of Endothelial Cell Growth Medium (Promocell, C-22010), Sertoli Cell Medium (Sciencell, 4521), and Leydig Cell Medium (Sciencell, 4511). Passaging was done using TrypLE Express Enzyme. 36 million cells were used for sorting with the Sertoli cell marker Thy-1 Cell Surface Antigen (THY1/CD90) using a PE-conjugated antibody for THY1/CD90 (R&D Systems, FAB2067P) and the EasySep™ Human PE Positive Selection Kit II (STEMCELL Technologies, 17664) with EasySep™ Magnet (STEMCELL Technologies, 18000), as per the manufacturer’s instructions, with the following specifications: 3 µg/mL FcR blocker, 3 µg/mL THY1/CD90 antibody, 75 µL/mL RapidSpheres™, 5 minutes incubation for separation 1, and 10 minutes incubation for separations 2 and 3. THY1/CD90+ cells were expanded on PLL-coated plates in Sertoli Cell Medium and passaged with TrypLE™ Express Enzyme.

### Immunocytochemistry

Cells were fixed for 15 minutes in 4% paraformaldehyde solution (PFA, Thermo Scientific, J19943-K2), permeabilized for 15 minutes in 0.1% Triton X-100 (Sigma, X100) in phosphate buffered saline (PBS), and blocked for 2 hours in 5% normal goat serum (NGS, Abcam, ab7481) in PBS. Primary antibodies were diluted in PBS as follows: anti-Hydroxy-Delta-5-Steroid Dehydrogenase, 3 Beta- And Steroid Delta-Isomerase 1 (HSD3β, Novus Biologicals, NB110-7844) 1:200, anti-Myosin Heavy Chain 11 (MYH11, Abcam, ab212660) 1:1000, anti-Octamer-Binding Protein 4 (OCT4, Abcam, ab184665) 1:500, anti-SRY-Box Transcription Factor 9 (SOX9, Abcam, ab76997) 1:500, anti-GATA Binding Protein 4 (GATA4, Abcam, ab84593) 1:100, anti-Nestin (Millipore, MAB5326) 1:200, anti-Thy-1 Cell Surface Antigen (THY1/CD90, Abcam, ab133350) 1:200, and incubated overnight at 4°C in the dark. Cells were rinsed 3 times with PBS for 15 minutes each at 4°C in the dark. Goat anti-Rabbit IgG (H+L) Highly Cross-Adsorbed Secondary Antibody Alexa Fluor 488 (Thermofisher, A-11034) or Goat anti-Mouse IgG (H+L) Highly Cross-Adsorbed Secondary Antibody Alexa Fluor 568 (Thermofisher, A-11031) were diluted 1:200 in PBS and incubated with the cells for 4 hours at 4°C in the dark. Cells were rinsed another 3 times with PBS for 15 minutes each at 4°C in the dark. 4′,6-diamidino-2-phenylindole (DAPI, Abcam, ab228549) was diluted to 2.5 µM in PBS and added to the cells for 15 minutes in the dark at room temperature, and then replaced by PBS. Cells were imaged using a Zeiss AXio Observer microscope equipped with laser excitation and fluorescence filters for AlexaFluor 488 and AlexaFluor 568 dyes, and ZEN Blue software. Image processing was done using ImageJ open source software.

### Real time quantitative polymerase chain reaction

Real time quantitative polymerase chain reaction (RT-qPCR) validation was done using PrimePCR™ Assays (Bio-Rad) with SYBR® chemistry. RNA was extracted using an RNeasy Plus Micro Kit (Qiagen, 74034), and checked for integrity using an Agilent 2200 Tapestation System with High Sensitivity RNA Screentape (Agilent, 5067-5579), High Sensitivity RNA ScreenTape Sample Buffer (Agilent, 5067-5580), and High Sensitivity RNA ScreenTape Ladder (Agilent, 5067-5581). cDNA was generated using iScript™ Reverse Transcription Supermix (Bio-Rad, 1708840) with a Tetrad2 Peltier Thermal Cycler (Bio-Rad). PrimePCR™ Primers (Bio-Rad) used were as listed in **Table 1**. RT-qPCR was done with SsoAdvanced™ Universal SYBR® Green Supermix (Bio-Rad, 1725270) on a LightCycler96 (Roche). Technical replicates were carried out in triplicate. Analyses was done in Excel and GraphPad Prism Software. Ct values were normalized to Glyceraldehyde-3-Phosphate Dehydrogenase (GAPDH). Outliers were detected using Grubbs’ Test, with α=0.05. Results of RT-qPCR are presented as the average Relative Quantification (RQ=2^-ΔΔCt^) values and the upper and lower RQ of the biological replicates. Any undetected samples were given a Ct value of the maximum number of cycles plus 1.

**Table 1.**
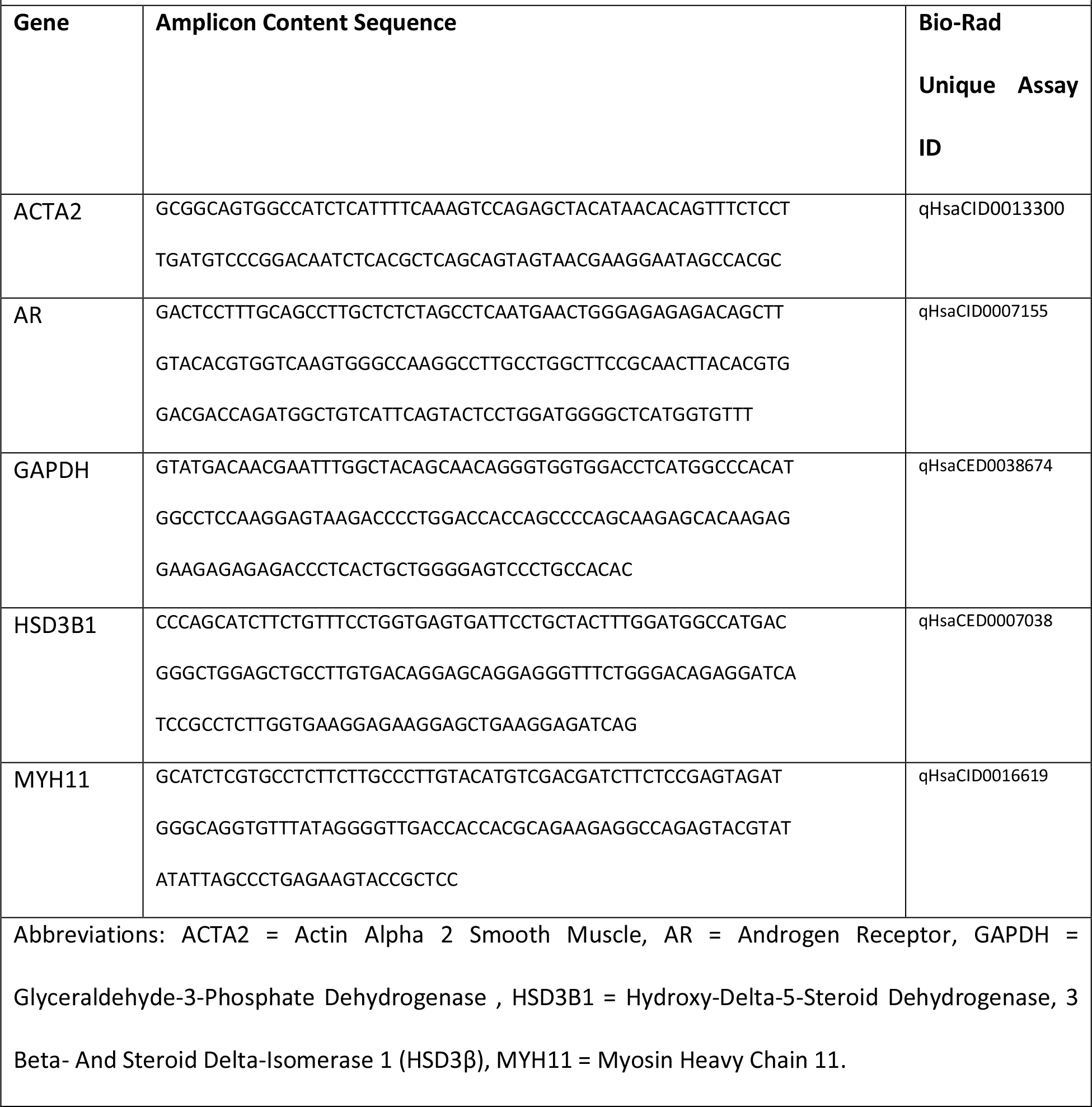
Amplicon content sequences and Unique Assay IDs for Bio-Rad PrimePCR™ primer pairs used in real time quantitative polymerase chain reaction (RT-qPCR) assays. Amplicon content sequences (amplicon sequence with additional base pairs added to the beginning and/or end of the sequence), and Unique Assay IDs for Bio-Rad PrimePCR™ primer pairs used for RT-qPCR.

### Statistics

Statistics were performed using GraphPad Prism software. Each experiment was performed in biological triplicate. For RT-qPCR results, significance was determined by comparing ΔCt values using a student’s unpaired two-tailed t-test, with α=0.05.

## RESULTS

To generate PTMs from hiPSCs (hPTMs), we employed a previously established Leydig cell differentiation protocol wherein hiPSCs acquire Leydig-like gene expression and steroidogenic function by way of patterning with growth factors and small molecule agonists of intracellular pathways (**Figure 2A**)(Chen, et al., 2020). These patterning factors were chosen based on their roles in Leydig cell differentiation *in vivo*, and validated by their ability to differentiate Leydig cells from SLCs *in vitro*, thus mirroring normal Leydig cell development from SLCs. To better mimic *in vivo* differentiation, we modified the protocol slightly by adding a preliminary step to first derive SLC precursor cells known as intermediate mesenchymal stem cells.(Wilhelm, et al., 2007) This step was hypothesized to improve the efficiency of differentiation by inhibiting early acquisition of ectodermal or endodermal lineages. We derived cells using this modified protocol, then substituted PDGFBB for PDGFAA to derive a second population of cells, and compared the resulting phenotypes in terms of gene expression and morphology. For a negative control, we isolated human primary Sertoli cells from a testis biopsy using MACS with an anti-THY1/CD90 antibody (**Figure 2B**). Sertoli cells are the spermatogenic regulatory cells of the niche. They share a cellular lineage with Leydig cells and PTMs, but have no steroidogenic or smooth muscle character, making them a suitable control for this study.

**Figure 2.**
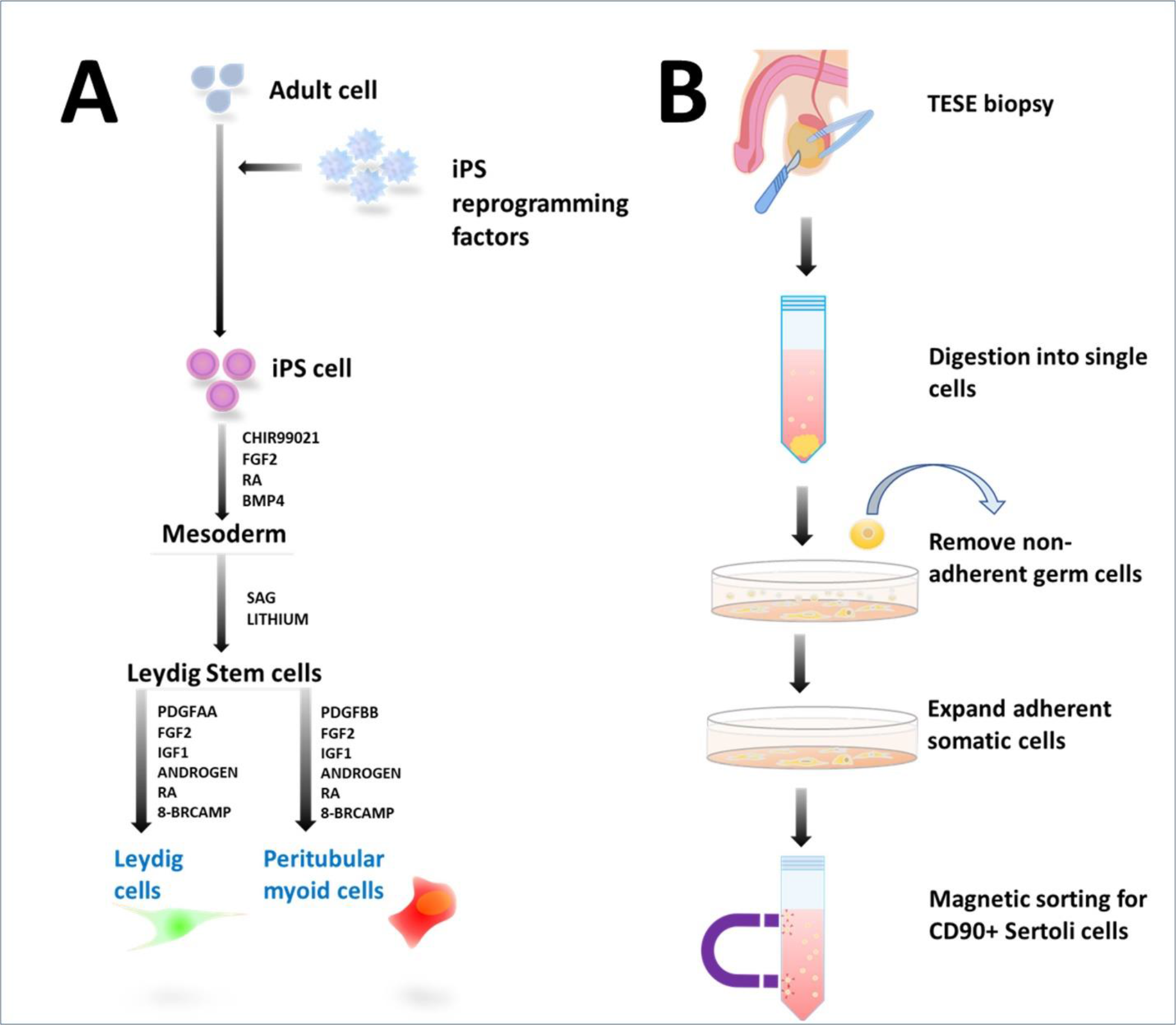
Experimental schematics for deriving Leydig cells and peritubular myoid cells from hiPSCs, and isolating primary Sertoli cells. **A)** Schematic representing the derivation of an intermediate mesoderm cell fate followed by either Leydig cell or peritubular myoid cell derivation. **B)** Schematic representing the isolation of primary Sertoli cells from a testicular biopsy by first digesting the biopsy into single cells, followed by removal of germ cells by differential plating, expansion of somatic cells and magnetic sorting. **Abbreviations**: iPS = induced pluripotent stem, FGF2 = Fibroblast Growth Factor 2, RA = retinoic acid, BMP4 = Bone Morphogenic Protein 4, SAG = smoothened agonist, PDGFAA = Platelet-Derived Growth Factor-AA, PDGFBB = Platelet-Derived Growth Factor-BB, IGF1 = Insulin-Like Growth Factor 1 = 8-BRCAMP = 8-bromo-cAMP, TESE = testicular sperm extraction, CD90 = /Thy-1 Cell Surface Antigen.

### Phenotype

In addition to being a known surface marker of cultured human Sertoli cells,(Chui, et al., 2011) THY1/CD90 has successfully been used to isolate rat SLCs using Fluorescent Activated Cell Sorting (FACS),(Guan, et al., 2019, Li, et al., 2016) therefore we expected sorted primary THY1/CD90 cell cultures to contain some contaminating SLCs in addition to Sertoli cells. However, the sorted cultures stained uniformly positive for the Sertoli cell-specific marker SOX9 in addition to THY1/CD90 (**Figure 2B-C**), while the Leydig cell marker HSD3β and peritubular myoid cell marker MYH11 were negative (**Figure 2A**), confirming a cell population highly enriched for Sertoli cells. The absence of THY1/CD90^+^ SLCs may be explained by differences between MACS and FACS techniques, such as the ability with FACS to further sort labeled cell populations based on size and granularity, or differences between rat and human SLC or Sertoli cell expression of THY1/CD90. Interestingly, we also observed Nestin expression in the sorted cells (**Figure 2B**). Since SOX9 was localized to nuclei, a sign of differentiated Sertoli cells,(Malki, et al., 2005) this was also unexpected. In addition to being associated with differentiating cells, Nestin is a well-known regulator of proliferation,(Bernal and Arranz, 2018) which is shown to resume in adult Sertoli cells under *in vitro* conditions.(Guo, et al., 2015) While Sertoli cells revert to a proliferative state *in vitro*, studies show that otherwise their global phenotypes remain stable, in agreement with the nuclear SOX9 localization noted in our cultures.(Guo, Hai, Yao, Chen, Hou, Li and He, 2015)

PDGFAA and PDGFBB-derived cells adopted immature Leydig cell-like and PTM-like phenotypes, respectively. PDGFAA-derived cells became spindle shaped whereas PDGFBB-derived cells acquired a flat polygonal shape, as expected based on previous descriptions of these cells in culture (**Figure 3A**).(Losinno, et al., 2012, Tung and Fritz, 1986) Moreover, spontaneous ring formation was noted in PDGFBB-derived cell cultures, a unique characteristic of purified PTMs in culture (**Figure 3A**).(Mishra, et al., 2012) Immunocytochemistry analysis revealed reactivity for GATA4, a testis-specific transcription factor(Ketola, et al., 2000, Viger, et al., 1998), in both cell types, confirming testis-specific lineages (**Figure 3A**). The Leydig cell steroidogenic enzyme HSD3β was present only in the PDGFAA-derived cells, while the peritubular myoid cell-specific smooth muscle protein MYH11 was only present in the PDGFBB-derived cells, confirming adoption of either Leydig cell-like or PTM-like phenotypes (**Figure 3A**).(Mayerhofer, 2013, Ye, et al., 2017) Nestin and CD90/THY1, two genes associated with SLCs,(Guan, Chen, Zhao, Hao, Chen, Ji, Wen, Lin, Ye and Chen, 2019, Ye, Li, Li, Chen and Ge, 2017) were also positive in both cell types (**Figure 3B**), suggesting that the cells were immature. Positive immunoreactivity for the Sertoli cell marker SOX9 was rare in both cell cultures (**Figure 3C**), denoting very little contamination by Sertoli-like cells. Likewise, immunoreactivity for OCT4, a pluripotency marker,(Takahashi, Tanabe, Ohnuki, Narita, Ichisaka, Tomoda and Yamanaka, 2007) was rare, indicating a very small presence of contaminating hiPSCs.

**Figure 3:**
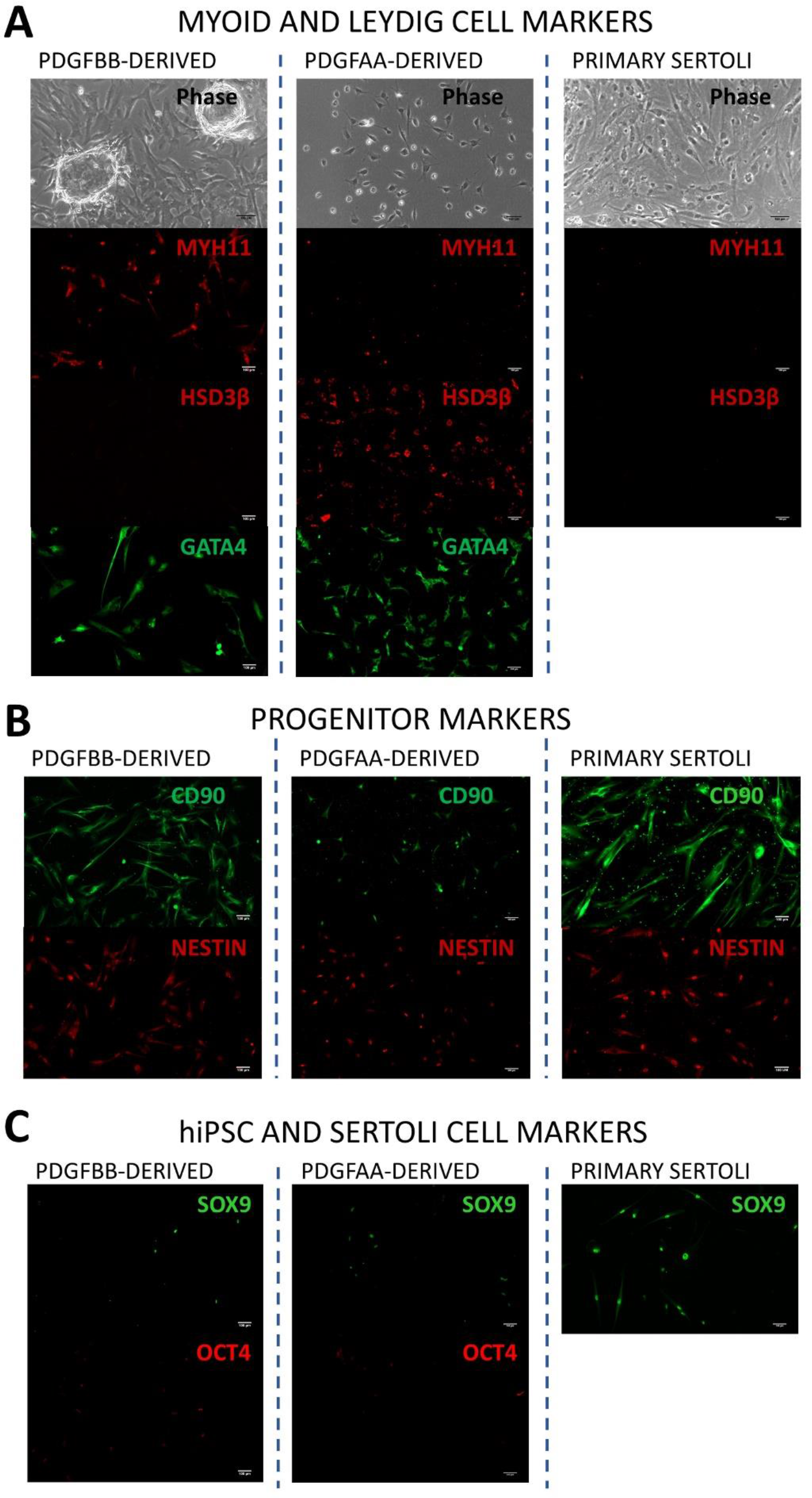
Phenotype characterization of PDGFAA-derived and PDGFBB-derived cells, and the primary Sertoli cell control. **A)** Peritubular myoid cell and Leydig cell phenotypes: phase contrast images and immunocytochemistry images of the myoid smooth muscle marker MYH11, the Leydig cell steroidogenic enzyme HSD3β, and the testicular somatic marker GATA4. **B)** Immature or differentiating phenotypes: the SLC and Sertoli cell marker CD90, and the differentiating cell marker Nestin. **C)** hiPSC and Sertoli cell phenotypes: the Sertoli cell marker SOX9, and the pluripotency marker OCT4. All scale bars are 100 µm. **Abbreviations:** PDGFBB = Platelet-Derived Growth Factor-BB, PDGFAA = Platelet-Derived Growth Factor-AA, MYH11 = Myosin Heavy Chain 11, HSD3B = Hydroxy-Delta-5-Steroid Dehydrogenase, 3 Beta- And Steroid Delta-Isomerase 1, GATA4 = GATA Binding Protein 4, CD90 = Thy-1 Cell Surface Antigen, hiPSC = human induced pluripotent stem cell, SOX9 = SRY-Box Transcription Factor 9.

### Gene expression

Gene expression analysis by RT-qPCR further confirmed distinct Leydig-like and PTM-like profiles in the PDGFAA and PDGFBB-derived cultures (**Figure 4A**). PDGFBB-derived cells upregulated expression of smooth muscle genes ACTA2 and MYH11 compared to PDGFAA-derived cells, by 23-fold [4-114] and 32-fold [6-161]. Compared to primary Sertoli cells, PDGFAA-derived cells upregulated MYH11 4-fold [1-11], whereas PDGFBB-derived cells upregulated MYH11 124-fold [26-623] (**Figure 4B**). In contrast to their immunocytochemistry profiles, no significant difference in HSD3β was observed between PDGFAA and PDGFBB-derived cells, suggesting differences in post-transcriptional regulation (**Figure 4B**). Nevertheless, compared to Sertoli cells, PDGFAA-derived cells increased gene expression of HSD3β 53-fold [15-190], verifying their steroidogenic phenotype (**Figure 4B**). Testicular cells rely upon androgen stimulation,(Mayer, et al., 2018, Meroni, et al., 2019, Wang, et al., 2009, Ye, Li, Li, Chen and Ge, 2017) therefore we examined their expression of androgen receptor. PDGFAA-derived cells possessed similar levels of androgen receptor gene expression as Sertoli cells (**Figure 4B**), while PDGFBB-derived cells exhibited greater expression by 3-fold [1-14] (**Figure 4B**), illustrating the adoption of androgen-dependent phenotypes by both cell types.

**Figure 4.**
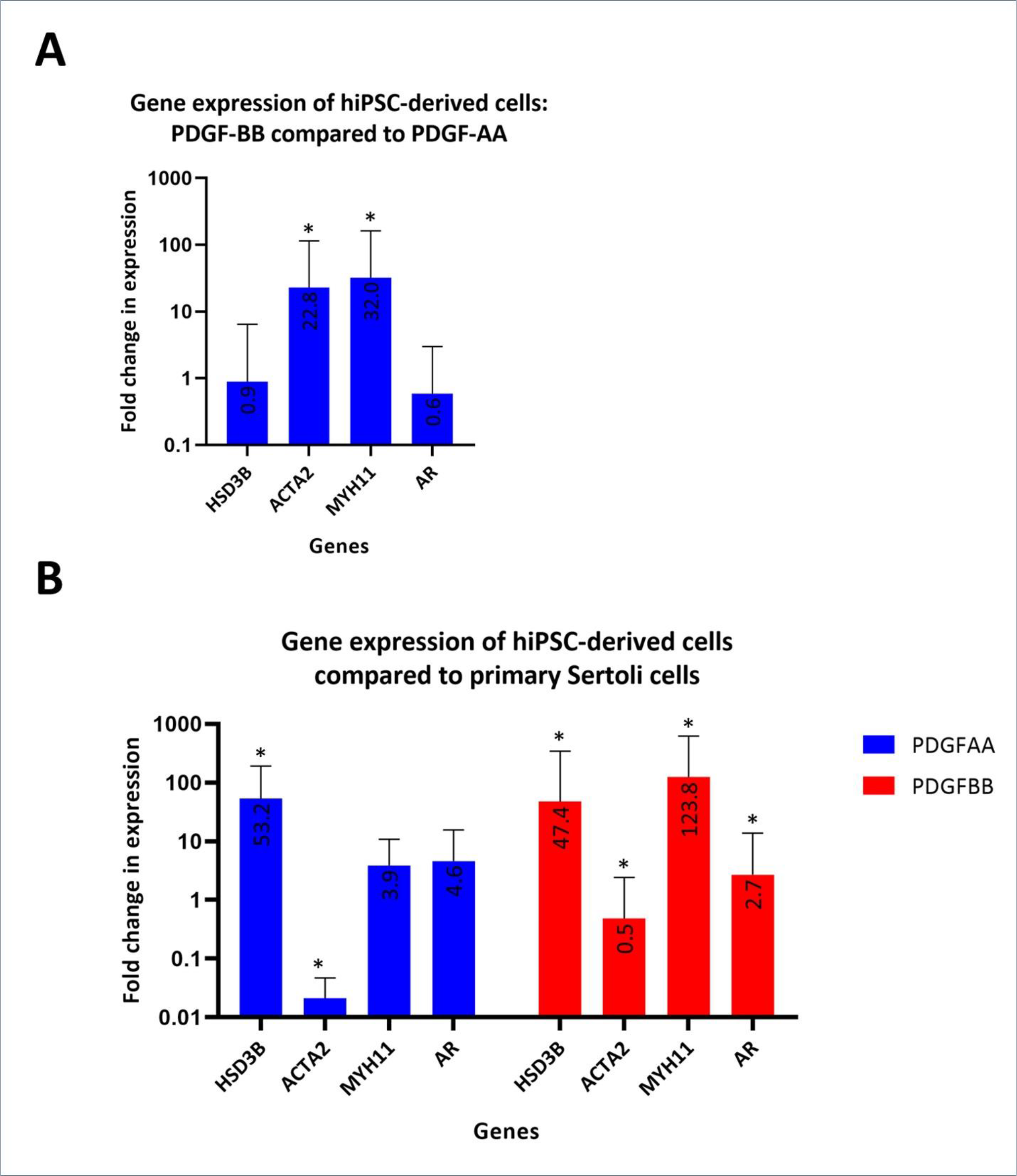
Gene expression of the PDGFBB and PDGFAA-derived cells. **A)** Fold change in gene expression in the PDGFBB-derived cells compared to the PDGFAA-derived cells. **B)** Fold change in gene expression in the PDGFBB-derived and PDGFAA-derived cells compared to the primary Sertoli cells. * indicates statistical significance. **Abbreviations:** HSD3B = Hydroxy-Delta-5-Steroid Dehydrogenase, 3 Beta- And Steroid Delta-Isomerase 1, ACTA2 = Actin Alpha 2, Smooth Muscle, MYH11 = Myosin Heavy Chain 11, AR = Androgen Receptor, PDGFAA = Platelet-Derived Growth Factor-AA, PDGFBB = Platelet-Derived Growth Factor-BB.

## DISCUSSION

In this study we derived PTM-like cells from hiPSCs and showed that the acquisition of their smooth muscle phenotype is dependent upon PDGFBB signaling. Furthermore, this derivation method mirrors what we know about *in vivo* patterning of Leydig and PTMs based on animal lineage tracing studies and animal knock-out models,(Gnessi, et al., 1995, Ye, Li, Li, Chen and Ge, 2017) suggesting its usefulness as a human developmental model. Indeed, the derivation of Leydig-like cells and PTM-like cells through the use of either PDGFAA or PDGFBB signaling using an otherwise identical patterning method highlights the closely related lineage between Leydig cells and PTMs, and is in agreement with recent single cell sequencing findings showing that human Leydig cells and PTMs arise from a shared progenitor.(Guo, Nie, Giebler, Mlcochova, Wang, Grow, DonorConnect, Kim, Tharmalingam, Matilionyte, Lindskog, Carrell, Mitchell, Goriely, Hotaling and Cairns, 2020) Furthermore, at the time of preparing this manuscript, another study was published showing that PDGFAA and PDGFBB delineate Leydig cell and PTM fates from the SLC progenitor stage in rat seminiferous tubule cultures.(Zhao, et al., 2021)

hPTM phenotypes were noted to possess immature gene expression of the differentiating marker Nestin and the progenitor marker CD90. Whether this was a result of missing microenvironmental factors necessary to mature PTMs or a consequence of PTM plasticity in response to unnatural *in vitro* culture conditions requires further investigation, but also highlights how little is currently understood regarding the effects of *in vitro* culture conditions on the PTM phenotype. Optimal human PTM culture conditions have not yet been explored due to their limited availability for research, with current models employing basic cell culture conditions including attachment onto 2-dimensional treated plastic surfaces, and growth in fetal bovine serum (FBS)-supplemented medium,(Albrecht, et al., 2006) leaving room for substantial improvements. In particular, 3-dimensional cultures are proving to be more useful model systems than 2-dimensional cultures by promoting biomimetic cell-cell and cell-matrix interactions, leading to more accurate data and greater insights.(Jensen and Teng, 2020) Indeed, a singular study of *in vitro* culture conditions for PTM culture showed that seminiferous tubule-derived extracellular matrix promoted an *in vivo*-like PTM histology whereas isolated components could not, suggesting that their correct histology depends upon a specific biomatrix.(Tung and Fritz, 1986) Another improvement to be explored could be replacement of FBS by a chemically defined or xeno-free medium for greater control over experimental variables, avoidance of batch-to-batch variability, and for ethical reasons.(Gstraunthaler, 2003, Gstraunthaler, et al., 2013, Jayme, et al., 1988, Witzeneder, et al., 2013) Because of its ill-defined composition of animal hormones, growth factors, and proteins, the use of FBS in current PTM models imparts unknowable and variable effects on the cells, limiting their experimental reproducibility and accuracy.

Animal and human studies have identified PTMs as critical to fertility and normal development, however differences in animal testicular development and cytoarchitecture(Guo, Nie, Giebler, Mlcochova, Wang, Grow, DonorConnect, Kim, Tharmalingam, Matilionyte, Lindskog, Carrell, Mitchell, Goriely, Hotaling and Cairns, 2020, Mayerhofer, 2013) suggest a need for complementary human models to gain further insight into human testicular development and causes of infertility. Until human primary peritubular myoid cells become more accessible to researchers, hPTMs could represent an alternative to bridge this gap in our understanding of their phenotype and functionality. While further characterization of hPTMs in terms of global gene expression profiling and functionality will better define the limitations, if any, with respect to how they can be used to model primary human PTMs, their potential to model some aspects of PTMs is readily apparent by their *in vivo*-like patterning, and their expression of smooth muscle and androgen receptor genes. For example, rodent PTM androgen receptor knock-out models show that Leydig cell differentiation and functionality is dependent on PTM androgen receptor-mediated activity,(Welsh, Moffat, Belling, de Franca, Segatelli, Saunders, Sharpe and Smith, 2012) while human studies have discovered a correlation between improperly differentiated Leydig cells and PTMs with testis cancer, Sertoli Cell Only (SCO) syndrome and Klinefelter syndrome.(Cina and Flannigan, 2020, Lottrup, et al., 2014, Winge, et al., 2018) The hPTM protocol presented here has the potential to model the relationship between PTMs and Leydig cells during their development.

The limitless supply of hPTMs also makes them a suitable cellular source for high throughput toxicity screening. The effect of environmental toxicants such as pesticides, dioxins, polychlorinated biphenyls, phthalates, alkylphenols and bisphenol A are known to alter reproductive function, and toxicology studies on rodents show that PTM development and function are highly susceptible.(Johnson, et al., 2007, Lara, et al., 2017, Saldutti, et al., 2013, Silvestroni, et al., 1999) Furthermore, patient-specific hiPSC-derived testis cells allows for non-invasive disease modelling for severe infertility conditions such as non-obstructive azoospermia. Now, with the addition of hPTMs to the previously described hiPSC-derived Leydig, Sertoli, and spermatogonial stem cells, 3-D modelling of spermatogenesis and the spermatogenic niche can be modelled and evaluated.

## CONCLUSION

Our novel differentiation method produces PTM-like cells as indicated by their morphology, protein expression, and gene expression. These cells will allow for ongoing understanding of the cellular interactions required for normal spermatogenesis in an *in vitro* setting. Furthermore, this method mirrors *in vivo* signaling necessary for the derivation of Leydig cell and PTM fates, opening the door for developmental studies.

## ACKNOWLEDGEMENTS

The authors would like to acknowledge the Vancouver Prostate Centre for their financial support, and the Lange Lab at the Vancouver Prostate Centre for their technical support.

## DECLARATIONS

### Ethics Approval

This study was performed in line with the principles of the Declaration of Helsinki. Approval was granted by the University of British Columbia Clinical Research Ethics Board (CREB approved protocol H18-03543). Experiments using hiPSCs in this study were not subject to ethics approval from the University of British Columbia Clinical Research Ethics Boards or Stem Cell Oversight Committee, since they were derived from somatic cells and not intended for transfer into humans or non-human animals.

### Consent to Participate

Human samples were obtained with informed consent and anonymized as per CREB protocol H18-03543.

### Availability of Data and Material

The data that support the findings of this study are available from the corresponding author upon reasonable request.

### Code Availability

Not applicable.

### Authors’ Contributions

Meghan Robinson: conceptualization, methodology, validation, formal analyses, investigation, writing – original draft, writing – review & editing, visualization. Luke Witherspoon: writing – original draft, writing – review & editing. Stephanie Willerth: resources, writing – review & editing, supervision. Ryan Flannigan: conceptualization, resources, writing – review & editing, supervision, project administration, funding acquisition.

